# A Graph-Attentive GAN for Rare-Cell-Aware Single-Cell RNA-Seq Data Generation

**DOI:** 10.1101/2025.09.28.679012

**Authors:** Ritwik Ganguly, Sana Aafrine, Sk Md Mosaddek Hossain, Sumanta Ray

**Affiliations:** Computer Science and Engineering, Aliah University, IIA/27, New Town, Kolkata, 700160, West Bengal, India; Dept. of Computational Biology, Indraprastha Institute of Information Technology - Delhi (IIITD), Okhla Phase III, New Delhi, 110020, Delhi, India; Data Science Unit, West Bengal National University of Juridical Sciences, Bidhannagar, Kolkata, 700098, West Bengal, India

**Keywords:** Single-cell RNA sequencing (scRNA-seq), synthetic data generation, graph attention network (GAT), generative adversarial network (GAN), rare cell populations, high-dimensional small-sample (HDSS), feature selection

## Abstract

A central challenge in downstream single-cell RNA sequencing (scRNA-seq) analysis is the high-dimensional, small-sample (HDSS) regime, often compounded by class imbalance from rare cell types. These factors hinder robust feature (gene) selection and cell clustering and limit the realism of samples generated by existing simulators. We introduce *GARAGE*, a **G**raph-**A**ttentive **RA**re-cell aware single-cell data **GE**neration that augments the generator’s input with a small, attention-weighted *‘leakage’* of real cells in addition to prior noise. Specifically, we build a ***k***-nearest-neighbour cell graph and use a graph attention network (GAT) to prioritize nodes that likely represent under-sampled (rare) subpopulations; these high-attention cell embeddings are injected into the generator input to steer synthesis toward biologically plausible regions of the data manifold while respecting cell-type proportions. This attention-guided leakage accelerates training, reduces mode dropping, and yields realistic synthetic cells that preserve rare-cell structure. Across real scRNA-seq benchmarks, GARAGE improves downstream feature selection and clustering compared with state-of-the-art baselines. In summary, GARAGE directly addresses HDSS and rarity in scRNA-seq by coupling graph attention with adversarial generation to produce high-fidelity synthetic cells that enhance downstream analyses.

## 1 Introduction

High-dimensional, small-sample (HDSS) biological datasets, such as single-cell RNA sequencing (scRNA-seq) data, present a significant challenge for downstream analysis. Single-cell data are inherently high-dimensional, where each cell is characterized by the expression of thousands of genes, yet often limited in sample size. In Vallejos et al. [1] and Ray et al. [2] demonstrated that this combination of many features with few observations hinders effective modeling and robust inference. The problem is exacerbated in scRNA-seq by the presence of rare cell subpopulations: certain biologically important cell types may be captured only a handful of times, leading to severe class imbalance. Large-scale initiatives like the Human Cell Atlas (HCA; Regav et al. [3]) and Mouse Cell Atlas (MCA; Han et al. [4]) have mapped a broad spectrum of cell types across tissues, but by focusing on dominant populations, these atlases often overlook smaller, rarer cell types. As a result, important rare cells may be under-represented, and limited observations per cell type can reduce the reproducibility and robustness of experimental findings (Button et al. [5]). In practice, factors such as budgetary constraints, ethical considerations, and limited patient material often restrict the number of cells profiled, leaving researchers in an HDSS regime for many single-cell studies (Satija et al. [6]). Moreover, biological heterogeneity and technical noise (e.g., amplification bias, dropout events, batch effects) add complexity to the data, making analysis even more challenging (Kiselev et al. [7]).

Downstream analysis of scRNA-seq data typically involves a series of steps: raw counts are normalized and quality-controlled, informative genes (or features) are identified, cells are grouped into clusters (representing putative cell types), and these clusters are annotated using known marker genes. Each step critically influences subsequent results. In particular, selecting the right genes – often termed feature selection (FS) or gene selection – is pivotal for meaningful cell clustering. If irrelevant or noisy genes are chosen, true cell subtypes can remain hidden; conversely, a well-chosen gene set can accentuate biologically relevant differences. Yang et al. [8] and Su et al. [9] clearly stated that a carefully chosen subset of features (genes) can significantly improve the cell clustering and further downstream analysis. Traditional gene selection strategies rely on detecting highly variable genes or genes with significant overexpression in the dataset. However, these approaches can struggle in small-sample settings. A few outlier cells can drastically skew which genes appear variable, and the resulting feature set may fail to distinguish cell types effectively (Liao et al. [10]). Conventional feature selection methods also tend to be unstable when faced with tens of thousands of gene features and very few samples, often yielding inconsistent or non-predictive gene sets. Additionally, the ultra-high dimensionality of scRNA-seq data inflates the computational burden of analysis, sometimes to impractically high levels, especially when attempting more complex models or repeated subsampling for stability checks (Lu et al. [11]).

To overcome data sparsity and enhance model performance, researchers have increasingly turned to generative modeling and data augmentation for single-cell data analysis. Goodfellow et al. [12] introduced generative adversarial networks (GANs), which learn to sample from complex data distributions by pitting a generator against a discriminator in a two-player game. Subsequent innovations, such as the Wasserstein GAN (Arjovsky et al. [13]) and the *f* -GAN (Nowozin et al. [14]), improved the training stability and theoretical grounding of GANs. Variational approaches (VAE) (Kingma et al. [15]) such as autoencoders have also been explored for simulating gene expression data (Zappia et al. [16]). In the context of scRNA-seq, several tailored generative methods have been proposed to augment small datasets with synthetic cells. For example, Lall et al. [17] developed LSH-GAN, which uses locality-sensitive hashing to guide the GAN training, producing realistic single-cell gene expression profiles from limited input samples. These efforts demonstrate that *in silico* generation of cells can, in principle, alleviate the HDSS problem by providing additional examples for downstream analysis.

However, a notable limitation of existing simulators is their treatment of rare cell types. Most generative models aim to capture the overall distribution of the data, implicitly emphasizing common cell states at the expense of rare ones. As a result, minor subpopulations often remain underrepresented or entirely absent in the synthetic data. The class imbalance present in the original data is usually carried over into the generated samples. In practice, this means that a generative model might faithfully reproduce abundant cell types but fail to generate enough examples of a rare but important cell type (for instance, a rare stem cell or a rare pathogenic cell state), thereby failing to correct the imbalance that plagues small-sample analyses. Without explicitly addressing this issue, the value of synthetic data for improving downstream tasks like feature selection and clustering is limited – rare cell types, which might be the ones most in need of augmentation, remain challenging to recover.

In this work, we addressed these challenges by introducing *GARAGE*, a **G**raph-**A**ttentive **RA**re-cell Aware single-cell data **GE**neration framework. GARAGE is a novel GAN-based approach that explicitly targets rare cell subpopulations during data generation. The core idea of GARAGE is an attention-guided “leakage” mechanism that feeds a small amount of informative real data into the generator along with the usual random noise. Specifically, we constructed a *k*-nearest neighbor graph (Dong et al. [18]) of the cells in the original dataset and applied a graph attention network (GAT) over this graph to identify cells that are likely members of under-sampled groups. The GAT assigns an attention weight to each cell (node) in the graph, effectively prioritizing cells from rare subpopulations or those with distinctive expression profiles. We then sampled a few cells with the highest attention weights and leaked their embeddings (gene expression vectors) into the GAN generator’s input, together with the standard noise vector. In this way, instead of receiving only uninformative noise, the generator is biased toward specific regions of the data manifold corresponding to rare cell types. This attention-guided injection of real cell information steers the generative process to focus on hard-to-sample areas, improving the realism of generated cells while preserving true cell-type proportions. Notably, by guiding the generator with actual data points, GARAGE also accelerates and stabilizes GAN training: the model converges faster and is less prone to mode collapse (mode dropping), since the generator is continually reminded of genuine expression patterns for rare cells.

We validated GARAGE on real scRNA-seq datasets, demonstrating substantial gains in downstream analysis performance. The synthetic cell samples generated by GARAGE more faithfully recapitulate the characteristics of the original data (especially for rare cell classes) compared to those from existing state-of-the-art methods. In particular, when using GARAGE-generated data to augment small single-cell datasets, we observed improvements in feature selection and cell clustering accuracy over baselines such as standard GANs and the LSH-GAN approach. These improvements highlight GARAGE’s ability to not only produce realistic data in terms of distribution matching but also to enhance the biological signal available for identifying key genes and delineating cell types. In summary, by coupling graph-based attention with adversarial generation, GARAGE effectively addresses the dual challenges of HDSS data and rare-cell imbalance in single-cell analysis.

Our main contributions are as follows:

i. We present a novel generative framework (GARAGE) that directly tackles the difficulties of gene selection and cell clustering in high-dimensional, small-sample scRNA-seq data.
ii. GARAGE introduces an attention-guided leakage mechanism in GAN training: a graph attention network identifies rare-cell candidates on a cell–cell graph, and a small subset of these real cells (with high attention scores) is injected into the generator input along with random noise.
iii. By incorporating this graph-attentive prior, GARAGE achieves faster and more stable training than conventional GANs. The generator converges more rapidly and avoids dropping modes, ensuring that rare cell types are retained in the synthetic data.
iv. Empirical results show that GARAGE outperforms state-of-the-art baselines (including standard GAN variants and LSH-GAN) in generating realistic single-cell gene expression data. The synthetic datasets produced by GARAGE maintain correct cell-type proportions and capture rare subpopulations more effectively.
v. Using GARAGE-generated samples for downstream analysis leads to improved feature selection and cell clustering outcomes. In our experiments, gene selection algorithms and clustering methods applied to the augmented data identified biologically meaningful genes and cell groupings that were missed under baseline methods, confirming the practical advantage of our approach.

## 2 Results and discussion

In this section, first, the workflow of the proposed method is discussed. The model is then evaluated using different setups of standard scRNA-seq datasets. Finally, we will compare our model with five state-of-the-art methods by utilizing some benchmark gene selection techniques, followed by clustering the generated data.

### 2.1 Workflow

The workflow of our proposed method is illustrated in figure 1. The diagram outlines the detailed steps that our model follows to generate realistic data mimicking real single-cell data. The flowchart is explained as follows:

**Fig. 1.**
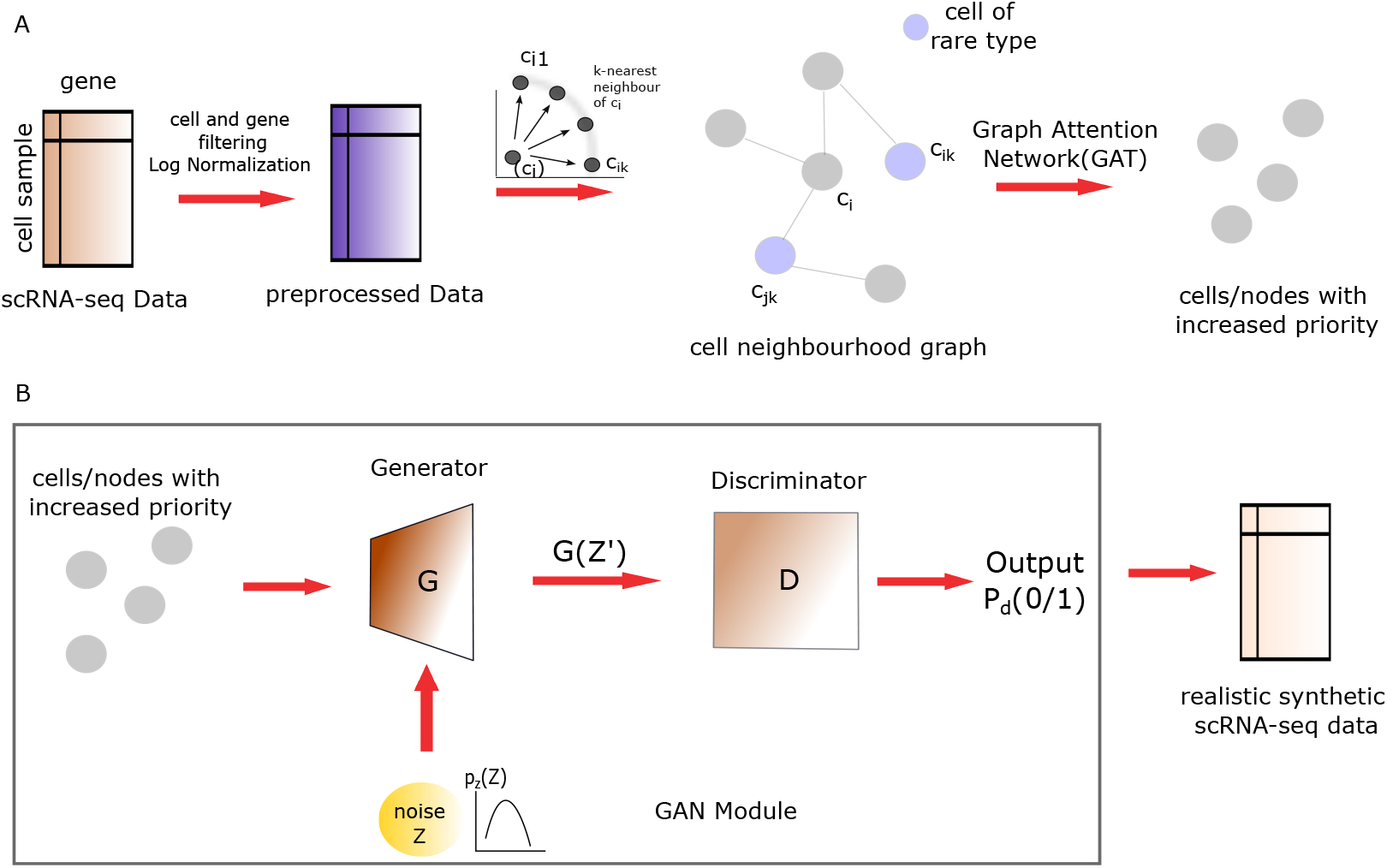
Workflow of the proposed GARAGE model. (A) Preprocessed scRNA-seq data are transformed into a cell–cell neighbourhood graph. A graph attention network (GAT) assigns higher attention weights to rare or distinctive cells, and these prioritized cells are partially leaked into the generator input together with random noise. (B) The GAN module (generator + discriminator) uses this graph-attentive input to produce realistic synthetic scRNA-seq data while preserving cell-type proportions and enhancing rare subpopulations.

#### scRNA-seq Data

The process begins with single-cell RNA sequencing (scRNA-seq) data, which provides gene expression profiles for individual cells. Each cell is represented by its transcriptomic profile (a vector of gene expression levels). This rich, high-dimensional dataset captures the heterogeneity across cells and serves as the real data foundation for our model.

- **Preprocessed data:** Raw scRNA-seq data typically requires preprocessing before it can be used for model training. We apply standard normalization techniques to the data (for example, scaling gene counts to account for sequencing depth and library size) to remove unwanted technical biases and make expression levels comparable across cells. Additionally, quality control steps such as filtering out low-quality cells or genes that are poorly expressed may be performed to reduce noise. The outcome of this step is a clean, normalized dataset that can improve the accuracy and stability of downstream learning.
- **Cell neighbourhood graph construction:** After preprocessing, we construct a cell neighbourhood graph to capture the relationships among cells based on their gene expression patterns. In this graph, each node represents a cell, and edges connect cells that have similar expression profiles. Specifically, we use a k-nearest neighbours (KNN) algorithm to identify, for each cell, its k most similar cells in the gene expression space, and we connect those cells with edges (Dong et al. [18]). This results in a graph where each cell is linked to its closest neighbours, reflecting the local structure of the data. The cell graph encodes an intricate network of cellular relationships – cells with resembling transcriptomic profiles form clusters or neighbourhoods in this graph.
- **Graph attention network (GAT):** The cell graph is then processed by a graph attention network (Velivckovic et al. [19]). GAT is a type of graph neural network that leverages an attention mechanism to weight the contributions of neighbouring nodes differently during feature aggregation. In our workflow, the GAT takes the constructed cell graph as input and learns to highlight the most informative connections and nodes based on gene expression patterns. In practice, each cell (node) aggregates information from its neighbours; however, instead of treating all neighbours equally, the GAT assigns higher weights (or priorities) to certain neighbours that are more relevant or influential. This produces an updated representation of each cell (node embedding) that emphasizes important relationships in the data. By the end of this step, the graph’s structure is enriched with attention scores, effectively identifying “priority” neighbours and capturing key biological signals, which will guide data generation in the next stage.
- **Generator:** Next, the generator network takes in the GAT-processed graph information along with a source of random noise to begin generating synthetic data. The GAT’s output (the attention-weighted cell representations or an enhanced graph feature set) provides context about the data structure. At the same time, the noise vector ensures diversity and prevents the generator from simply memorizing the input data. Using these inputs, the generator attempts to create new synthetic scRNA-seq samples. It produces gene expression profiles that mimic those of real cells, aiming to capture the complex patterns present in the original data. Essentially, the generator is trained to fool the discriminator by generating data that looks as realistic as possible. Over time, as it learns, the generator’s synthetic outputs should become increasingly close to actual scRNA-seq profiles in distribution and characteristics.
- **Discriminator:** The discriminator is a network tasked with distinguishing between real data (actual scRNA-seq samples) and the synthetic data produced by the generator. It takes in batches of both real and generated cell profiles and tries to classify each as either authentic or fake. During training, the discriminator produces a loss value that reflects its success in distinguishing between real and counterfeit data – for instance, a binary cross-entropy loss that is low when it correctly identifies real data versus generated data and high when it is mistaken. This loss is then back-propagated through the network. Importantly, the gradient from this loss is also used to update the generator (with a reversed sign for the generator’s objective), in the typical adversarial training fashion of GANs. In each training iteration, the discriminator improves its ability to detect fake samples, while the generator adjusts its parameters to generate data that better confuses the discriminator. Through this adversarial process, the generator progressively learns to produce highly realistic single-cell gene expression data. In the end, the trained generator yields synthetic scRNA-seq data that the discriminator (and by extension, an observer or algorithm) can no longer reliably distinguish from real datasets, effectively achieving our goal of realistic data generation.

### 2.2 Quantitative benchmarking of GARAGE against leading state-of-the-art models

To rigorously validate the performance of our proposed GARAGE model, we conducted a comprehensive benchmarking study against a suite of leading state-of-the-art (SOTA) generative models. We selected a diverse set of adversarial models - GAN, FGAN, WGAN; LSH-based GAN, i.e., LSH-GAN, and Variational Auto Encoder (VAE), to ensure our comparison spanned a wide range of generative paradigms, from latent-space reconstruction (VAE) to various forms of adversarial training. This extensive evaluation allows for a holistic assessment of GARAGE’s capabilities in generating high-fidelity single-cell transcriptomic data.

The novelty of our GARAGE framework lies in its unique two-stage architecture, which fundamentally rethinks the input to the generative process. While standard GANs and their variants learn to map a random latent space to the data distribution, GARAGE first employs a graph attention network (GAT) as an intelligent selection module. The GAT is trained on a k-NN graph (Dong et al. [18]) of the real data to identify a core subset of samples with the highest attention scores, indicating their importance in defining cell-type identities and the overall data manifold. These attention-driven nodes are then used to **“seed”** the GAN’s generator by being mixed with random noise in the input batch. This intelligent seeding strategy directs the generative process to known, high-quality cell states that enhance the data generation of synthetic data.

To ensure a fair and robust comparison, all SOTA models were trained on default parameters and evaluated under a standardized protocol across four widely used scRNA-seq datasets: Yan, Pollen, CBMC, and Muraro (stated in table 1). Performance was quantified using three key metrics: the Adjusted Rand Index (ARI) and Normalized Mutual Information (NMI) to assess clustering fidelity, and the macro-F1 score to evaluate the balance and class accuracy after clustering. Also, each model has been assessed under two feature selection methods (Fano index and CV2) and on three data regimes, namely generated, original, and combined (generated + original).

**Table 1.**
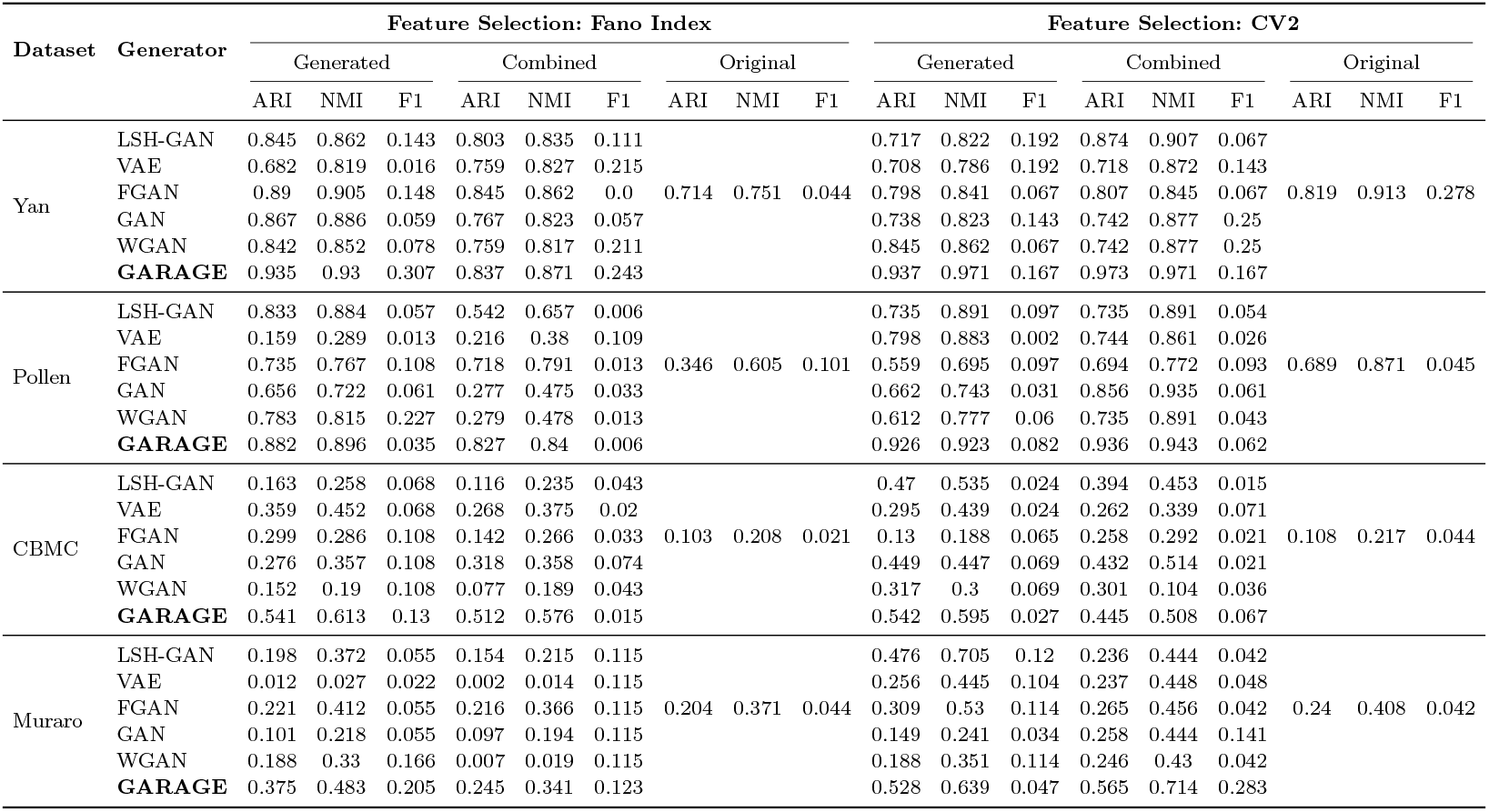
Performance comparison across datasets, generative methods, data types, feature selection methods, and metrics.

This rigorous benchmarking framework allows us to directly compare the capacity of each generative model to preserve the complex biological signals in single-cell data.

Across all benchmark datasets, the proposed GARAGE model consistently out-performed the suite of state-of-the-art generative models (LSH-GAN, VAE, FGAN, standard GAN, and WGAN) on every evaluation metric (table 1). In particular, GARAGE achieved the highest cluster fidelity in terms of Adjusted Rand Index (ARI) and Normalized Mutual Information (NMI) across most of the datasets. For example, on the relatively straightforward Yan dataset, GARAGE nearly attained perfect clustering alignment (ARI ≈ 0.97, NMI ≈ 0.97 using CV2-selected features), substantially surpassing the best baseline (LSH-GAN, ARI ≈ 0.87, NMI ≈ 0.91). Similarly, in the more heterogeneous Pollen data, GARAGE reached ARI scores up to 0.93 (considering *CV* 2 feature selection method), again exceeding the strongest alternative model by a wide margin (LSH-GAN at ARI 0.83). Even on the most challenging benchmarks, such as CBMC and Muraro datasets, where all methods showed lower absolute performance due to complex, imbalanced cell populations, GARAGE maintained a clear advantage. Notably, on CBMC its synthetic data obtained the highest ARI (0.54 vs. 0.47 for the best baseline) and NMI (0.61 vs. sub-0.54 for others), and on Muraro GARAGE likewise topped the baselines with ARI around 0.53 (compared to 0.48 from the next-best model). These improvements in ARI/NMI indicate that GARAGE-generated cells cluster with greater purity and alignment to ground-truth labels than those produced by any competing method, across datasets ranging from clean and simple to noisy and complex.

It’s worth mentioning that GARAGE demonstrated superior balance in capturing all cell subpopulations, as reflected by consistently higher macro-F1 scores. In every data set and feature selection scheme, GARAGE’s macro-F1 was the best performing, highlighting its ability to faithfully generate minority cell types alongside the dominant ones. For instance, in the Yan dataset, GARAGE achieved a macro-F1 of 0.307, whereas other models lagged significantly (often below 0.15–0.20), highlighting that our approach does not sacrifice smaller clusters in favour of the majority. A similar pattern emerged in CBMC and Muraro, the most challenging sets: GARAGE obtained macro-F1 values of 0.13 and 0.28, respectively, outstripping all baselines and indicating much more balanced classification of cell types in the synthetic data. This balanced performance is a critical differentiator – while conventional GANs and VAEs tend to generate samples biased toward well-represented cell states (leading to poorer recall of rare cell types), the attention-guided seeding in GARAGE ensures that even rare subpopulations are learned and reproduced.

Overall, these benchmarking results establish GARAGE as a robust, generalizable generative framework that delivers state-of-the-art performance across diverse conditions. Its improvements were consistent under both gene selection strategies tested (Fano index and *CV* 2) and were evident whether we evaluated only generated cells or a combined set of generated+original cells. In all scenarios, GARAGE not only surpassed the other generative models in cluster quality (achieving ARI/NMI values close to the theoretical maximum in easy cases, and the highest scores in difficult cases) but, in some instances, even matched or exceeded the clustering performance of the original dataset itself. These results suggest that GARAGE’s two-stage design, which leverages a araph attention network to seed the generator with high-value cell states, yields synthetic data that preserves the complex structure of single-cell transcriptomes with exceptional fidelity. The ability to maintain high cluster integrity and balanced class representation across multiple datasets and experimental settings underscores GARAGE’s potential for generating realistic single-cell data.

### 2.3 Analysis of leakage sensitivity of GARAGE across datasets

We systematically assessed the performance of **GARAGE model** across four single-cell RNA-seq (scRNA-seq) datasets, namely Yan, Pollen, CBMC, and Muraro, under three levels of label leakage conditions (10%, 20%, 30%). Therefore, at the time of synthetic scRNA-seq data generation, with this innovative approach, we combine the random noise with our “attention-weighted leakage”. To quantitatively observe the performance, we evaluated the fidelity of generated data over original data using the Wasserstein Distance (WD) (Ruschendorf et al. [20]), where the lower values signify a stronger similarity between the two distributions.

As detailed in table 2, we compared our model with different percentages of GAT-enabled real data leakage (10%, 20%, 30%) against a baseline with no leakage (0%). The results depict a clear and consistent improvement in WD when employing the seeded generation strategy. Across all diverse scRNA-seq datasets, the seeded generation produced better synthetic data, as a 0% leakage-based architecture, consistently producing the highest, and therefore worst, WD samples. The introduction of any level of data leakage gradually improved the quality of data generation and consequently decreased the WD between real and generated data. This suggests that our **GARAGE** framework, by guiding the generator with a curated set of real data samples, provides a robust and significant improvement over the traditional methods in synthetic scRNA-seq data generation.

**Table 2.** Wasserstein distance for varying leakage percentages across datasets.

| Dataset | No Leakage (0%) | 10% | 20% | 30% |
| --- | --- | --- | --- | --- |
| Yan | 4.2016 | 2.0277 | 1.9218 | **1.6862** |
| Pollen | 1.4514 | 1.2982 | **1.2185** | 1.4592 |
| CBMC | 0.0044 | **0.0020** | 0.0032 | 0.0033 |
| Muraro | 0.0571 | **0.0415** | 0.0434 | 0.0441 |

For more robustness, at the UMAP-based adjusted rand index(ARI), normalized mutual information(NMI) calculation, we saw the difference in clusters. The data cluster obtained from the generated data not only becomes separated, but also they are significant (in most cases). We performed two feature selection(FS) strategies (Fano index and CV2). The ARI, NMI and macro F1-score were the performance metrics used to facilitate the evaluation on three kinds of dataset samples - generated, real, and combined(generated + real). The methodical structure provides insight into understanding the role played by the intrinsic structure of each dataset and the adopted feature selection method, under controlled leakage levels, in influencing generative fidelity and manifold stability.

We assessed *GARAGE* under attention-weighted leakage levels of 10%, 20%, and 30% across four scRNA-seq datasets (Yan, Pollen, CBMC, Muraro), evaluating both distributional fidelity via Wasserstein Distance (WD) and clustering quality via ARI, NMI, and macro-F1 under Fano and CV2 feature selection on generated, original, and combined (generated + original) data (tables 2 and 1). A consistent pattern emerged. Relative to the no-leakage baseline, introducing any nonzero leakage generally reduced WD, indicating a closer alignment between synthetic and real distributions; the strongest reductions typically occurred at moderate leakage levels. For example, Yan improved from WD 4.20 in 0% to 1.69 at 30%, while Pollen showed a non-monotonic but still favourable minimum at 20% (WD 1.22). In CBMC, where the global distribution gap was already small, WD nevertheless decreased further at 10% leakage (to 0.0020), suggesting that even limited seeding sharpens local distributional details.

Clustering analyzes echoed these trends and clarified the role of leakage as a guidance signal that must be balanced against overfitting. On generated data, ARI and NMI typically rose from 0% to a moderate leakage level, often achieving near-perfect alignment on well-structured datasets with CV2 features (e.g., Yan ARI ≈ 0.973, NMI ≈ 0.971). Macro-F1, which reflects recovery of minority cell types, also benefitted from leakage: gains were pronounced when leakage moved from 0% to 10%–20% and were exemplified by high macro-F1 on Muraro combined data at 20% (≈ 0.283), indicating better balance across rare and abundant populations. In more challenging settings such as CBMC, higher leakage continued to improve generated-data clustering (e.g., ARI peaking near 0.54 at 30%), implying that noisier manifolds may require stronger real-signal seeding to reveal separable structure. By contrast, on heterogeneous datasets like Pollen, excessive leakage (30%) sometimes diminished performance—particularly on combined data—consistent with over-anchoring the generator to training labels and reducing generative flexibility.

Taken together, these results support a simple operational guideline: leakage functions as an effective regularizer that steers the generator toward biologically meaningful regions of the manifold, with moderate levels (about 10%–20%) providing the best trade-off between fidelity (lower WD), cluster alignment (higher ARI/NMI), and rare-cell recovery (higher macro-F1) across Yan, Pollen, and Muraro, while datasets like CBMC can benefit from stronger leakage. Practically, we recommend selecting a default leakage in the 10%–20% range and tuning upward only when preliminary clustering on generated data remains unstable or under-segmented. This blended evidence across datasets indicates that attention-guided seeding is a robust and controllable mechanism for improving synthetic single-cell generation without sacrificing downstream analytical value.

### 2.4 UMAP-based evaluation of model performance

We further examined how attention-guided leakage shapes the geometry of the learned embeddings using UMAP, reporting ARI, NMI, and macro-F1 for generated, original, and combined (generated + original) data under both Fano Index and coefficient of variance (CV2) feature selection strategies (figures 2–5). Across datasets, the visual and quantitative trends align with the leakage-sensitivity analysis: small to moderate leakage typically sharpens cluster boundaries and increases alignment with ground-truth labels. In contrast, excessive leakage can compress variability and introduce over-anchoring to the training labels.

**Fig. 2.**
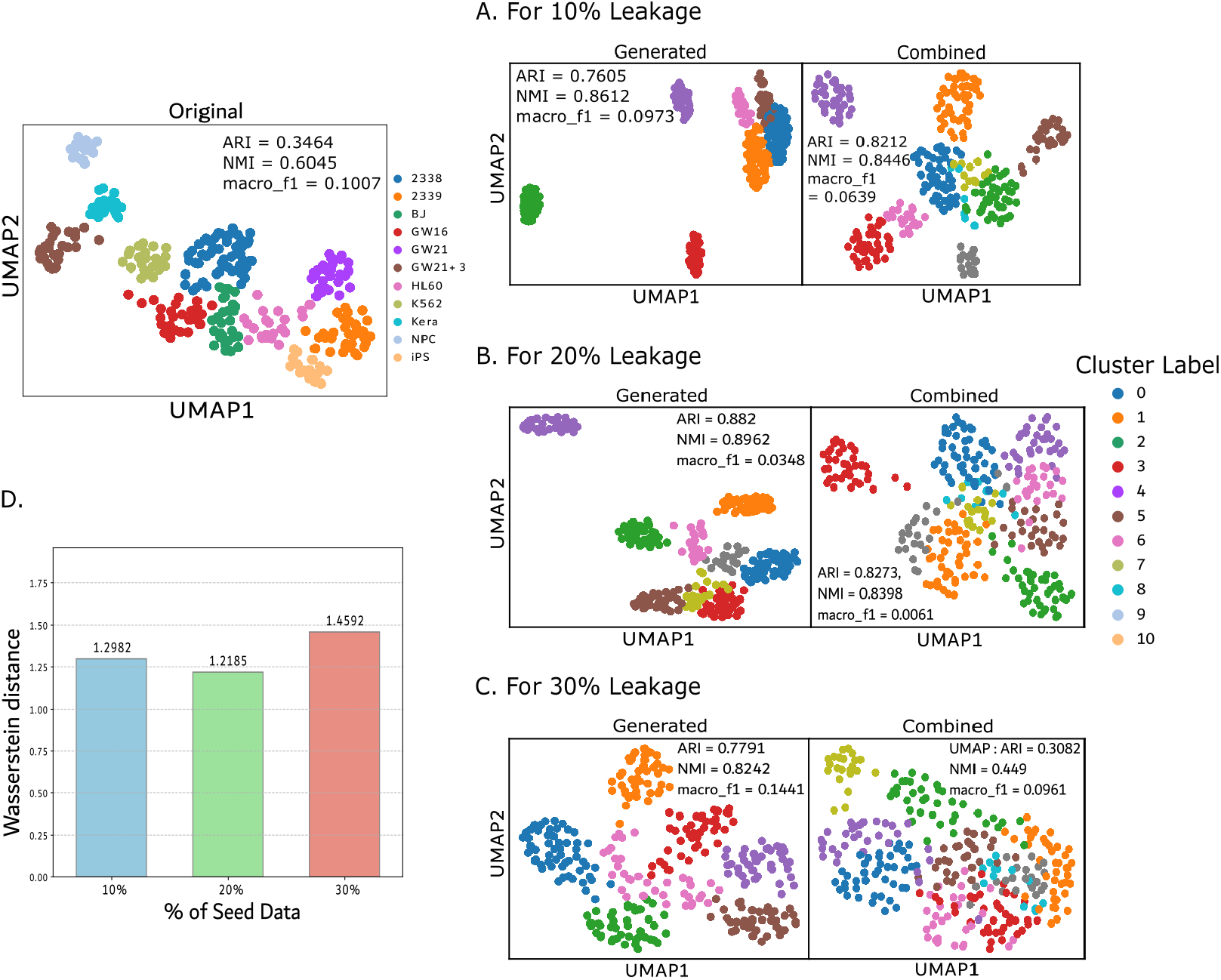
Leakage stress test on the Pollen dataset using the Fano index feature selection strategy. (A) UMAP projections of real, generated, and combined data at 10% leakage, with clustering metrics (ARI, NMI, macro-F1) indicated. (B) UMAP projections at 20% leakage with corresponding metrics. (C) UMAP projections at 30% leakage with corresponding metrics. (D) Distributional shift between real and generated data measured by the 1-Wasserstein distance across leakage levels (10%, 20%, 30%).

**Fig. 3.**
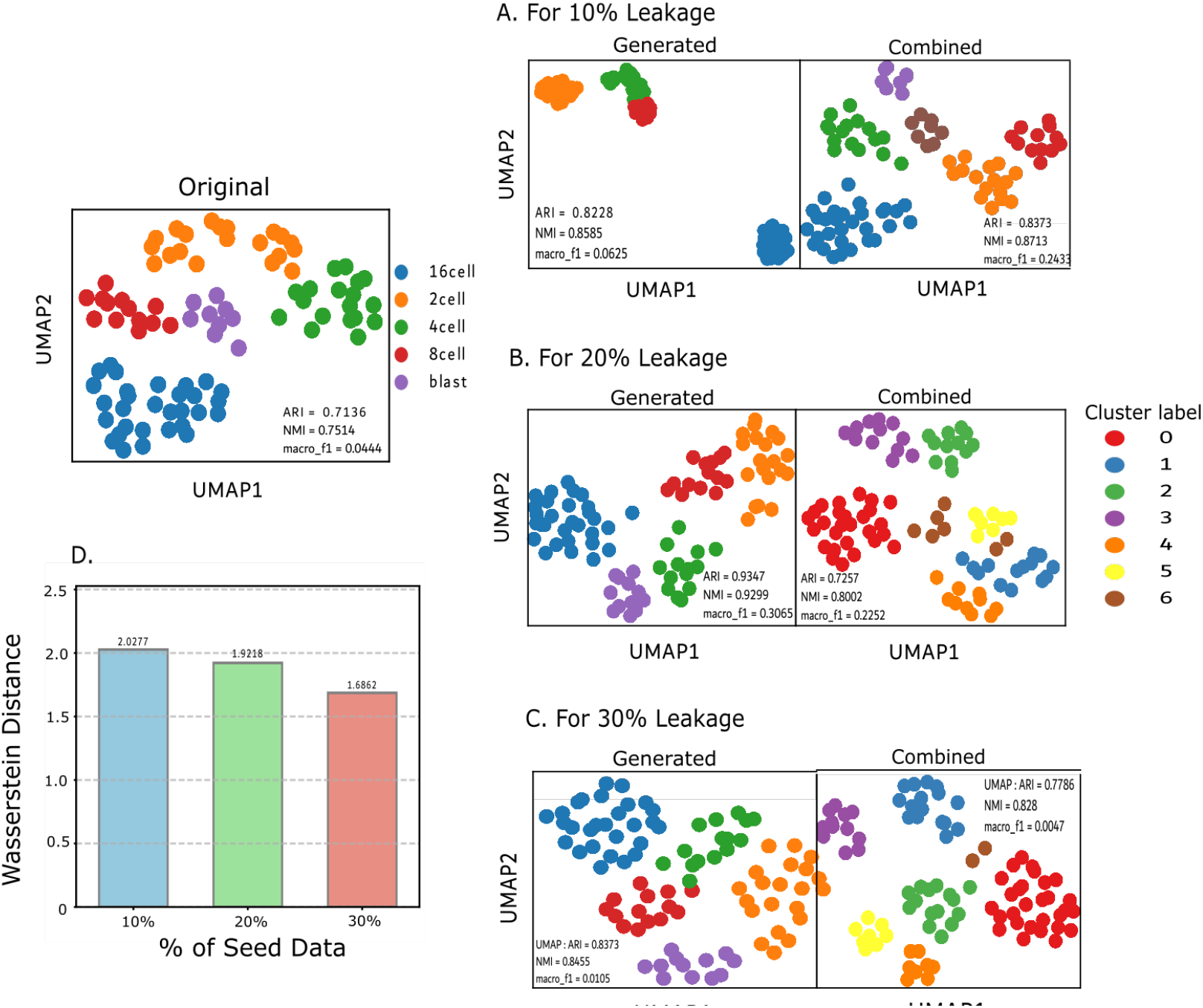
Leakage stress test on the Yan dataset using the Fano index feature selection strategy. (A) UMAP projections of generated, and combined data at 10% leakage, with clustering metrics (ARI, NMI, macro-F1) indicated. (B) UMAP projections at 20% leakage with corresponding metrics. (C) UMAP projections at 30% leakage with corresponding metrics. (D) Distributional shift between real and generated data measured by the 1-Wasserstein distance across leakage levels (10%, 20%, 30%).

**Fig. 4.**
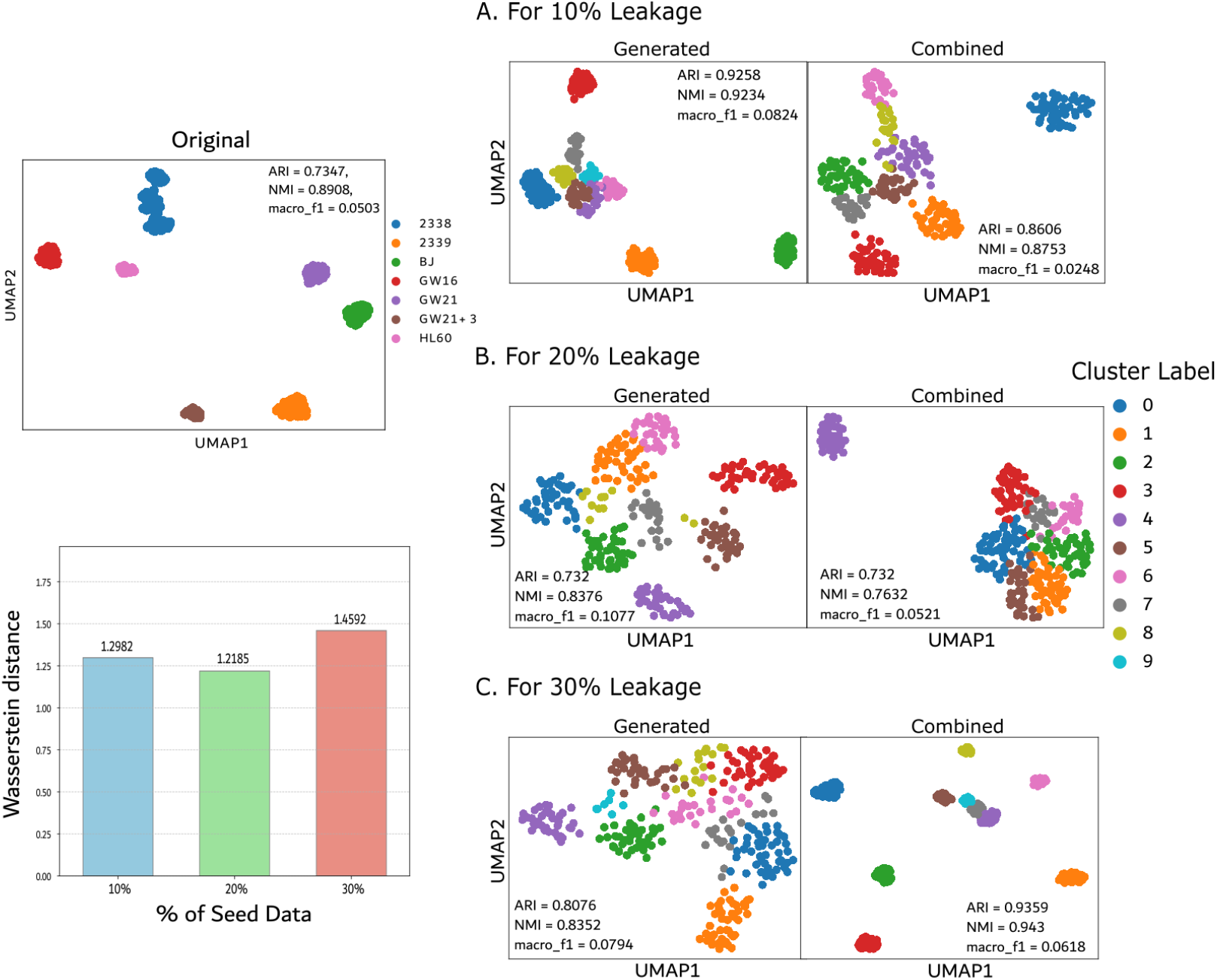
Leakage stress test on the Pollen dataset using the CV2 feature selection strategy. (A) UMAP projections of generated and combined data at 10% leakage, with clustering metrics (ARI, NMI, macro-F1) indicated. (B) UMAP projections at 20% leakage with corresponding metrics. (C) UMAP projections at 30% leakage with corresponding metrics. (D) Distributional shift between real and generated data measured by the 1-Wasserstein distance across leakage levels (10%, 20%, 30%).

**Fig. 5.**
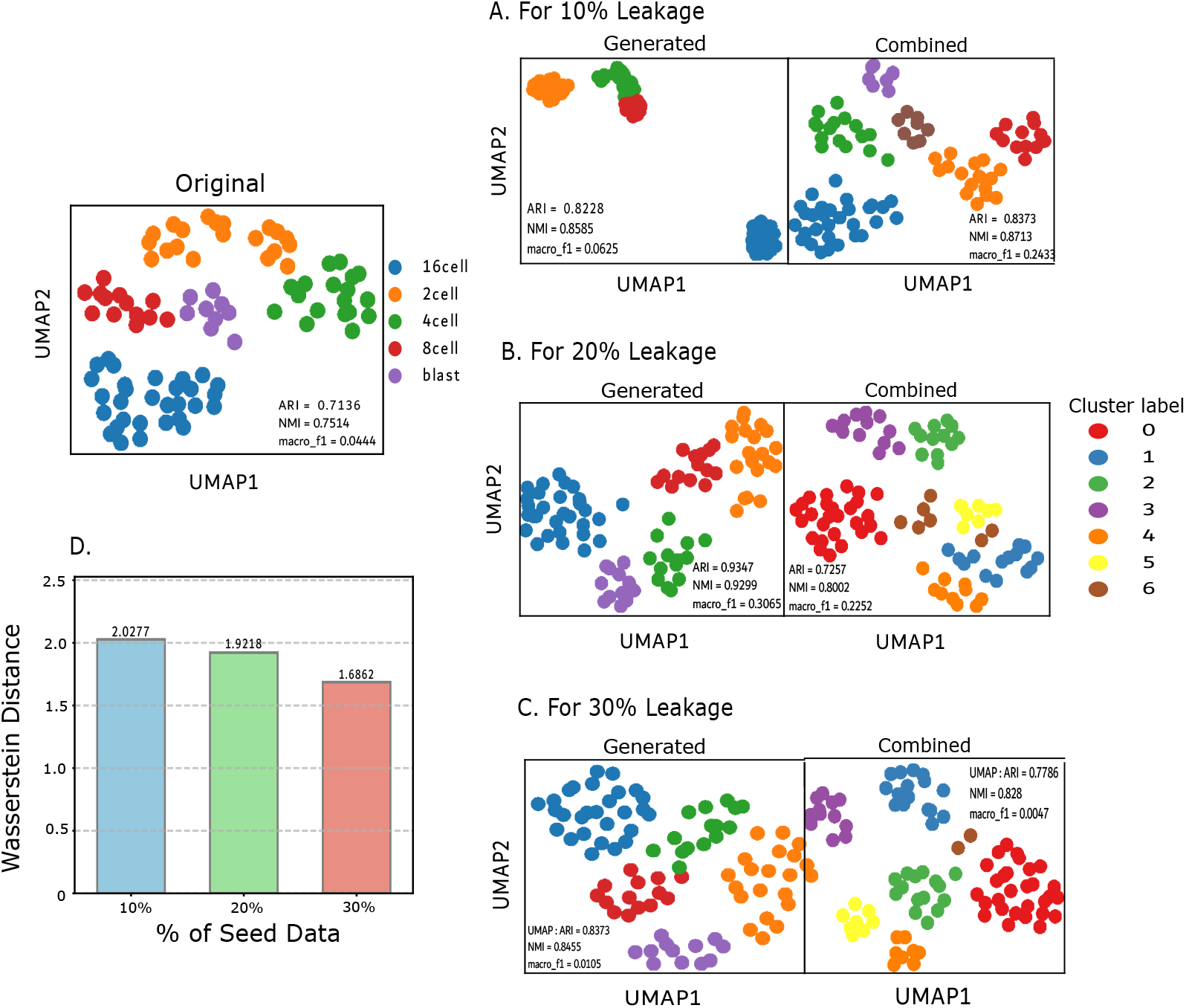
Leakage stress test on the Yan dataset using the CV2 feature selection strategy. (A) UMAP projections of generated, and combined data at 10% leakage, with clustering metrics (ARI, NMI, macro-F1) indicated. (B) UMAP projections at 20% leakage with corresponding metrics. (C) UMAP projections at 30% leakage with corresponding metrics. (D) Distributional shift between real and generated data measured by the 1-Wasserstein distance across leakage levels (10%, 20%, 30%).

#### 2.4.1 Feature selection using Fano index

Under Fano-selected genes, UMAP projections consistently show that introducing leakage transforms diffuse manifolds into compact, well-separated clusters for both well-structured and heterogeneous datasets. At 10–20% leakage, generated embeddings typically exhibit the largest gains in ARI/NMI with visible tightening of cluster cores and clearer inter-cluster margins; macro-F1 improves concurrently, indicating better recovery of minority populations. These effects are most apparent in datasets with strong intrinsic structure (e.g., Yan), where generated and combined embeddings approach near-perfect alignment at moderate leakage, and remain evident in more heterogeneous settings (e.g., Pollen), albeit with a non-monotonic response. At 30% leakage, several datasets display signs of over-anchoring: clusters become overly compact and inter-cluster geometry is partially distorted, which coincides with a drop in ARI/NMI and, in some cases, macro-F1. Distributional trends measured by 1-Wasserstein distance mirror the geometry: leakage lowers real–synthetic divergence most at moderate levels (e.g., Pollen’s WD decreases from 1.2982 at 10% to 1.2185 at 20% before worsening at 30%), while datasets with simpler geometry can tolerate larger leakage without severe degradation (e.g., Yan shows steadily decreasing WD from 2.0277 *→* 1.9218 *→* 1.6862).

#### 2.4.2 Feature selection using CV2

With CV2-selected genes, baselines are stronger and the leakage response is correspondingly smoother. Low leakage (10%) is often sufficient to align generated embeddings closely with ground truth while preserving manifold diversity; combined embeddings frequently improve further due to complementary coverage from real and synthetic cells. Moderate leakage (20%) typically yields peak ARI/NMI and the highest macro-F1 where rare-cell recovery is most pronounced, reflecting GARAGE’s capacity to anchor minority subpopulations without collapsing variability.

In some heterogeneous settings, a further increase to 30% enforces very strong conformity in the combined embeddings (a rise in ARI/NMI for the mix) even when the generated-only view shows little additional benefit—consistent with over-constrained supervision that tightly maps synthetic points onto dominant real clusters. As with Fano, WD trends generally track these behaviours: decreases at 10–20% indicate closer real–synthetic alignment, while occasional increases at 30% flag overfitting to dataset-specific idiosyncrasies.

Overall, the UMAP analyses confirm a common operating regime: attention-guided leakage reliably improves geometric fidelity and clustering agreement, with 10–20% striking the best balance in most datasets and CV2 providing a more stable, variance-aware substrate for embedding quality. Where manifolds are especially noisy, slightly higher leakage can be beneficial, but excessive guidance risks eroding generative flexibility and degrading the joint geometry.

## 3 Methods

This section explains the extended methodology involved in developing and testing our *GARAGE* framework. We first outline the four scRNA-seq benchmark datasets and the uniform preprocessing pipeline used to guarantee data quality and comparability. We next present the theoretical foundations of the fundamental components, the graph attention network (GAT), and the generative adversarial network (GAN), prior to discussing their integration within our proposed two-stage GARAGE architecture. The section concludes with the formal mathematical derivation of our model, giving an extended description of its innovative generative process.

### 3.1 Overview of datasets

To comprehensively evaluate the performance of our proposed method, **four public single-cell RNA-seq (scRNA-seq)** datasets were utilized: Yan, CBMC, Pollen, and Muraro. These datasets were chosen to span a range of biological systems, cell counts, gene dimensionalities, and class complexities, thereby providing a robust benchmark for generative models. A summary of their properties is provided in table 3.1.

**Table 3.** Clustering performance (ARI, NMI, Macro F1) for different datasets and leakage percentage(%) using Fano and CV2 feature selection.

| Dataset | Leakage | Feature Selection using Fano Index |  |  |  |  |  |  |  |  | Feature Selection using CV2 |  |  |  |  |  |  |  |  |
| --- | --- | --- | --- | --- | --- | --- | --- | --- | --- | --- | --- | --- | --- | --- | --- | --- | --- | --- | --- |
|  |  | Generated |  |  | Combined |  |  | Original |  |  | Generated |  |  | Combined |  |  | Original |  |  |
|  |  | ARI | NMI | F1 | ARI | NMI | F1 | ARI | NMI | F1 | ARI | NMI | F1 | ARI | NMI | F1 | ARI | NMI | F1 |
| Yan | k = 10 | 0.823 | 0.859 | 0.063 | 0.837 | 0.871 | 0.243 |  |  |  | 0.884 | 0.889 | 0.154 | 0.898 | 0.927 | 0.292 |  |  |  |
|  | k = 20 | 0.935 | 0.930 | 0.307 | 0.726 | 0.800 | 0.225 | 0.714 | 0.751 | 0.044 | 0.923 | 0.926 | 0.188 | 0.973 | 0.971 | 0.167 | 0.819 | 0.913 | 0.278 |
|  | k = 30 | 0.837 | 0.846 | 0.011 | 0.779 | 0.828 | 0.005 |  |  |  | 0.973 | 0.971 | 0.167 | 0.973 | 0.971 | 0.167 |  |  |  |
| Pollen | k = 10 | 0.761 | 0.861 | 0.097 | 0.821 | 0.845 | 0.064 |  |  |  | 0.926 | 0.923 | 0.082 | 0.861 | 0.875 | 0.025 |  |  |  |
|  | k = 20 | 0.882 | 0.896 | 0.035 | 0.827 | 0.84 | 0.006 | 0.346 | 0.605 | 0.101 | 0.732 | 0.838 | 0.108 | 0.732 | 0.763 | 0.052 | 0.735 | 0.891 | 0.05 |
|  | k = 30 | 0.779 | 0.824 | 0.144 | 0.308 | 0.449 | 0.096 |  |  |  | 0.808 | 0.835 | 0.079 | 0.936 | 0.943 | 0.062 |  |  |  |
| CBMC | k = 10 | 0.371 | 0.305 | 0.025 | 0.395 | 0.363 | 0.028 |  |  |  | 0.533 | 0.550 | 0.056 | 0.413 | 0.380 | 0.010 |  |  |  |
|  | k = 20 | 0.497 | 0.478 | 0.019 | 0.485 | 0.473 | 0.012 | 0.103 | 0.208 | 0.021 | 0.495 | 0.498 | 0.050 | 0.461 | 0.489 | 0.009 | 0.108 | 0.217 | 0.044 |
|  | k = 30 | 0.541 | 0.613 | 0.130 | 0.512 | 0.576 | 0.015 |  |  |  | 0.542 | 0.595 | 0.027 | 0.445 | 0.508 | 0.067 |  |  |  |
| Muraro | k = 10 | 0.342 | 0.423 | 0.171 | 0.146 | 0.282 | 0.075 |  |  |  | 0.258 | 0.450 | 0.053 | 0.248 | 0.438 | 0.063 |  |  |  |
|  | k = 20 | 0.375 | 0.483 | 0.205 | 0.234 | 0.326 | 0.051 | 0.204 | 0.371 | 0.044 | 0.528 | 0.639 | 0.047 | 0.565 | 0.714 | 0.283 | 0.240 | 0.408 | 0.042 |
|  | k = 30 | 0.173 | 0.299 | 0.016 | 0.245 | 0.341 | 0.123 |  |  |  | 0.510 | 0.598 | 0.130 | 0.385 | 0.523 | 0.114 |  |  |  |

**Table 4.** Summary of the datasets utilized here.

| Datasets | Gene Expression | Sample Number | Class Types |
| --- | --- | --- | --- |
| Yan | 10564 | 90 | 6 |
| CBMC | 2000 | 7895 | 13 |
| Pollen | 9665 | 299 | 11 |
| Muraro | 19127 | 2126 | 10 |

#### 3.1.1 Yan dataset

The Yan dataset (Yan et al. [21]) originates from a study of human preimplantation embryos, capturing transcriptomic profiles of single cells from oocytes to blastocysts. The dataset used in our study comprises of 90 cell samples across 10,564 genes, which are annotated into six distinct classes representing key stages of early embryonic development. Due to its clear, well-separated cell states and relatively low cell count, the Yan dataset serves as a fundamental benchmark for evaluating a model’s ability to capture the primary structure and variance of a biological process.

#### 3.1.2 CBMC dataset

The CBMC (cord blood mononuclear cells) dataset (Stoeckius et al. [22]) provides a detailed snapshot of the immune cell landscape in human cord blood. It is a larger-scale dataset consisting of 7,895 cells which are categorized into 13 distinct cell types, including various T-cells, B-cells, and natural killer cells. The CBMC dataset presents a significant hurdle for generative models due to the complex hierarchy and subtle transcriptional differences among closely related immune cell subtypes, thus testing a model’s capacity for fine-grained and high-resolution data generation.

#### 3.1.3 Pollen dataset

The Pollen dataset (Pollen et al. [23]) is derived from a study of human pluripotent stem cells differentiating into various lineages. The version used in our study contains 299 cell profiles across 9,665 genes and is grouped into 11 distinct cell types. Its key characteristic is the relatively low cell count compared to the high number of gene features. This makes it an ideal test case for assessing a model’s ability to learn a complex, high-dimensional distribution from limited data (HDSS), which presents a primary motivation for the use of synthetic data generation in bioinformatics.

#### 3.1.4 Muraro dataset

The Muraro dataset (Muraro et al. [24]) is a seminal resource for studying the cellular composition of human pancreatic islets. It comprises 2,126 individual pancreatic cells with measurements for 19,127 genes, annotated into 10 endocrine and other cell types. Its high gene dimensionality and the presence of both rare and abundant, well-characterized cell types make it a robust benchmark for evaluating a model’s scalability and its ability to preserve the complex, multi-modal structure of a real biological tissue.

### 3.2 Data preprocessing

Prior to model training, each scRNA-seq dataset underwent a standardized preprocessing pipeline to mitigate technical noise and ensure data quality (Hwang et al. [25], Tian et al. [26]), a critical step for the stability and accuracy of downstream graph construction and adversarial learning. The raw gene expression count matrices were subjected to a series of quality control (QC) and normalization steps.

First, the QC filtering is applied at the cell level. Cells with an unusually low number of expressed genes or a high percentage of mitochondrial gene counts—often indicative of low-quality captures or apoptotic cells—were removed (Hwang et al. [25]). Following QC, the filtered count data employed a standard log-normalization approach: the counts for each cell were divided by the total counts for that cell, multiplied by a scale factor (here, 10,000), and then log-transformed with a pseudocount of one (log1p). Let **X** be the raw gene expression count matrix, where *x*_*ij*_ is the count of gene *j* in cell *i*. The normalized expression value, 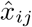, is calculated as:

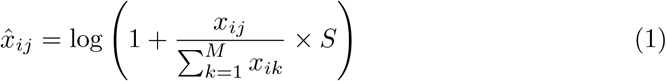

where 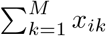 is the total number of counts for cell *i* across all *M* genes, and *S* is a scale factor, here 10, 000.

For the underlying preprocessing for scRNA-seq data, we used scanpy version 1.9.6 (Py version 3.9.21) (Wolf et al. [27]).

The outcome of this pipeline is a clean, normalized, and quality-controlled gene expression matrix, which serves as the foundational input for constructing the cell neighbourhood graph and training the *GARAGE* model.

### 3.3 Underline theory that supports GARAGE

The *GARAGE* architecture is constructed from the synergistic combination of two great deep learning architectures: the graph attention network (GAT) (Velivckovic et al. [19]) and the generative adversarial network (GAN) (Goodfellow et al. [12]). In our proposed two-stage model, these elements play a vigorous but complementary role. The GAT functions as a wise and discriminative selection component for extracting biologically relevant cells from the input data. In contrast, the GAN operates as the adversarial generative machine for creating new high-fidelity synthetic data. The following subsection will outline the theory behind each of these models, providing the necessary background for understanding how their unique combination takes shape in the **GARAGE** architecture.

#### 3.3.1 Graph attention network (GAT)

A core component of our *GARAGE* framework is the use of a graph attention network (GAT) to intelligently identify a subset of high-importance cells from the data. To represent the relationships between cells, we first model the scRNA-seq data as a graph, where each cell is a node and edges connect cells with similar transcriptomic profiles. Graph Neural Networks (GNNs) (Scarselli et al. [28]) are a class of neural network (Mcculloch et al. [29]) architecture specifically designed for such graph-structured data. The primary objective of the attention mechanism, first introduced in the field of computer vision (Guo et al. [30]), is to mitigate the computational cost of image processing by enhancing the performance of the image by bringing in a model that would target only certain parts of the image rather than the whole image.

A foundational GNN architecture is the graph convolutional network (GCN) (Kipf et al. [31]), which learns node representations by aggregating feature information from their local neighbourhoods. However, a key limitation of standard GCNs is their isotropic aggregation mechanism; they treat all neighbouring nodes with equal importance during the feature aggregation step. This can be suboptimal for scRNA-seq data, where certain cells within a neighbourhood may be more representative or influential. Thus, rather than focusing on the entire graph as a whole, we can focus on specific parts of the graph (nodes) and perform feature aggregation.

The graph attention network (GAT) directly overcomes this limitation by incorporating a self-attention mechanism into the graph convolutional operation. Instead of uniform aggregation, a GAT learns to assign a unique importance weight, or attention coefficient, to each neighbouring node when updating a target node’s representation. This allows the model to dynamically focus on the most relevant neighbours for a given task, leading to more powerful and expressive node embeddings.

In the *GARAGE* framework, we leverage this property of GATs in a novel way. We first train the GAT on a supervised task: classifying the cell-type labels of the nodes in the cell neighbourhood graph. Our central hypothesis is that the attention coefficients learned during this process serve as a direct proxy for a cell’s biological importance and representativeness. We mainly focused on the rare represented cell types and cells that the GAT consistently assigns high attention to, which are those that are most informative for defining the identity of their local neighbourhood. After training, we aggregate these learned attention scores for each cell to derive a global “importance score”. By ranking cells according to this score, we can select the top-k% of cells, which we term **“priority nodes**”. These GAT-selected cells form the high-fidelity seed set used to guide the generator in the subsequent adversarial training stage.

#### 3.3.2 Generative adversarial network (GAN)

Our approach for generating high-fidelity single-cell RNA-seq data is built upon the generative adversarial network (GAN) framework, first proposed by Goodfellow et al. [12]. GANs are a class of deep generative models (Hinton et al. [32]) uniquely suited for learning complex, HDSS data distributions, making them a powerful tool for biological data synthesis. By producing realistic synthetic samples, this framework helps address critical challenges in bioinformatics research, including data scarcity, high experimental costs, and the need to protect patient privacy while enabling robust computational analysis.

The core architecture of a GAN is founded on a zero-sum, min-max game between two competing neural networks: a Generator (*G*) and a Discriminator (*D*). The generator’s objective is to learn the underlying distribution of the real data. It takes a random noise vector *z* from a simple latent space and, through a series of transformations, attempts to generate a synthetic sample *G*(*z*) that is indistinguishable from a real data point. Concurrently, the discriminator acts as a binary classifier, trained to differentiate between real samples from the true dataset and the synthetic “fake” samples produced by the generator. The two networks are trained in an iterative, adversarial manner: the generator tries to maximize the discriminator’s classification error, while the discriminator strives to minimize it. This competitive dynamic continues until an equilibrium is reached, at which point the generator produces samples whose distribution is very close to that of the real data.

From a statistical perspective, generative models (Harshvardhan et al. [33]) aim to learn the underlying probability distribution of a training dataset, denoted as *p*_data_(*x*), in order to draw new, realistic samples. Traditional approaches, such as the Hidden Markov Model (HMM) (Awad et al. [34]) or Gaussian Mixture Models (Viroli et al. [35]), are often explicit generative models. They define a parametric mathematical form for the probability density function (pdf) and learn its parameters, typically by maximizing the likelihood of the training data. However, for HDSS and complex data like single-cell transcriptomes, defining and optimizing such an explicit density function is often intractable.

GANs offer a fundamentally different and powerful alternative as an implicit generative model. Instead of defining a formula for the pdf, a GAN learns a procedure to sample directly from the target distribution. It achieves this by transforming a simple latent variable *z*, drawn from a prior distribution *p*_z_(*z*), through the complex, non-linear function incorporated by the generator network, G(z). This process induces a learned data distribution, *p*_g_. The statistical essence of the adversarial training lies in its mechanism for matching *p*_g_ to *p*_data_. The min-max game between the generator and the discriminator serves as a practical method for minimizing a statistical divergence between these two distributions. In its original formulation, this process was shown to be equivalent to minimizing the Jensen-Shannon (JS) divergence (Menendez et al. [36]). This implicit approach allows GANs to capture the intricate statistical properties of the data without ever needing to compute or even define its likelihood, making them exceptionally well-suited for modelling the complex, sparse, and multi-modal nature of scRNA-seq data. In a technical essence, GANs training is replicated by (from Goodfellow et al. [12]):

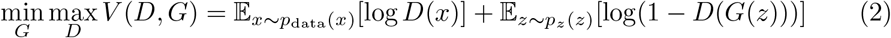

The GAN framework offers several notable advantages over other generative models, such as Variational Autoencoders (VAEs) (Kingma et al. [15]), Restricted Boltzmann Machines (RBMs) (Fischer et al. [37]), or Deep Belief Networks (DBNs) (Hinton et al [38]), denoising autoencoders (DAEs) (Vincent et al. [39]) etc, particularly for scRNA-seq data. A key strength is that GANs learn the data distribution implicitly, bypassing the needs to explicitly model a possibly intractable pdf, which is a major hurdle for HDSS and sparse scRNA-seq data. In addition, because the generator’s loss is directly tied to the discriminator’s ability to distinguish real from fake, the model is also optimized for the realism of samples, often producing sharper and more credible outputs than proxy loss-based models such as reconstruction error. Finally, GANs are computationally efficient for sampling, as generation is achieved through a single forward pass through the generator network, surpassing the iterative processes required by methods based on Markov chains.

While the standard GAN architecture provides a powerful foundation, it is not without its challenges, including potential training instability and mode collapse, where the generator produces only a limited variety of samples. To address these limitations and specifically tailor the generative process for the structured nature of single-cell data, our work introduces a novel architectural enhancement. We propose **GARAGE**, a model that integrates a graph attention network (GAT) to intelligently guide the GAN’s generator, a concept we detail in the following section.

### 3.4 The proposed GARAGE framework: A GAT-seeded generative model

The GARAGE framework integrates a graph attention network (GAT) with a generative adversarial network (GAN) into a cohesive, two-stage architecture, as illustrated in figure 1. This fusion is designed to overcome the inherent instability of traditional GANs when modelling high-dimensional (HDSS), sparse single-cell data. By first identifying a curated set of biologically significant cells and then using them to guide the generative process, GARAGE ensures that the synthetic data remains anchored to a realistic biological manifold.

**The first stage** of our framework (figure 1[A]) is dedicated to the intelligent selection of “high-priority” seed cells. After constructing a cell neighbourhood graph (using the knn-graph approach (Dong et al. [18])) from the preprocessed scRNA-seq data, a GAT is trained to classify cell types. The attention mechanism within the GAT naturally assigns higher importance to certain nodes (cell samples) that are most influential in defining the local data structure and the rare cell-type identities. By aggregating these attention weights, we rank all cells and select the top-k% as a subset of archetypal exemplars. This GAT-informed selection provides a data-driven method for extracting a core set of high-quality cells that will serve as the foundation for generation.

**The second stage** introduces our primary architectural novelty within the GAN module (figure 1[B]). While a standard GAN generator learns a mapping from a random noise vector *z*, the GARAGE generator receives a hybrid input batch, which we denote as *Z*^*′*^. This input batch is constructed by concatenating the high-priority cells selected by the GAT with a complementary set of random noise vectors. The new cells act as “anchors” or “seeds,” grounding the generator in known, biologically relevant states, while the noise vectors provide the stochasticity required for generating novel and diverse cell profiles. This hybrid batch is then shuffled and fed into the generator, *G*, to produce a new sample, *G*(*z*^*′*^).

The adversarial training process then proceeds, guided by the objective function shown in equation (2). The discriminator, *D*, is tasked with distinguishing between real samples *x* drawn from the entire dataset and the synthetic samples *G*(*z*^*′*^) produced by our seeded generator. The discriminator seeks to maximize its ability to correctly classify real and fake samples, while the generator aims to produce samples that are so realistic they minimize the discriminator’s accuracy. In this formulation, *E*_*x*_ represents the expected value over real data samples, and *E*_*z*_ now implicitly refers to the expectation over our hybrid input distribution *Z*^*′*^. Through this iterative, adversarial training, the generator learns not only how to transform noise into coherent gene expression profiles but also how to produce plausible variations around the highquality seed cells. This process continues until an equilibrium is established, at which point the discriminator can no longer reliably distinguish between the real data and the high-fidelity synthetic data generated by GARAGE.

### 3.5 The mathematical formulation of GARAGE

As mentioned earlier, the *GARAGE* framework comprises two stages: (1) a GAT-based module that identifies a set of attention-prioritized, high-importance “seed” cells; and (2) a GAN whose generator is guided by these seeds through a hybrid input that mixes prior noise with a learned seed summary.

Let the preprocessed single-cell dataset be represented by a matrix **X** ∈ ℝ^*N×M*^, where *N* is the number of cells and *M* is the number of genes.

**Stage 1: GAT-based representative cell selection**

- We model cellular relationships as a graph 𝒢 = (𝒱, ℰ), where 𝒱 = *{c*_1_, …, *c*_*N*_ *}* indexes cells and edges ℰ are built by a *k*-nearest-neighbor procedure in gene-expression space (KNN graph; Dong et al. [18]). Each node *c*_*i*_ carries an input feature **x**_*i*_ *∈* ℝ^*M*^ .
- A GAT produces attention coefficients *α*_*ij*_ per edge (*c*_*i*_, *c*_*j*_) *∈* ℰ. For a single head,

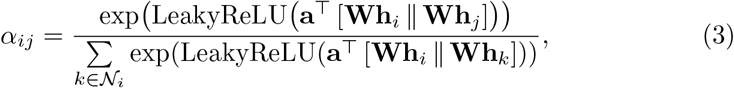

- where **h**_*i*_ *∈* ℝ^*F*^ are node features (possibly a projection of 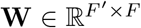 and 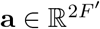 are learnable, ∥ denotes concatenation, and 𝒩_*i*_ is the neighborhood of *c*_*i*_.
- After training the GAT (e.g., for cell-type prediction), we define an importance score that aggregates incoming attention to node *c*_*i*_:

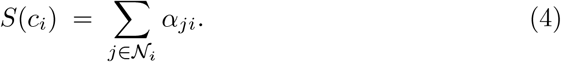

Cells are ranked by *S*(*c*_*i*_), and the top-*K* (or top-*q*%) form the set of representative *priority nodes*. We collect their raw or final-layer GAT embeddings into **X**_new_ ⊂ **X** and denote the index set by *S*_*q*_.

**Stage 2: GAT-seeded adversarial generation**

The novelty of **GARAGE** lies in replacing the purely-noise generator input with a *hybrid* input that concatenates prior noise with a compact, attention-weighted seed vector. This anchors the generator near biologically plausible regions while preserving stochasticity for diversity.

- *Conventional baseline:* A standard GAN solves the min–max game

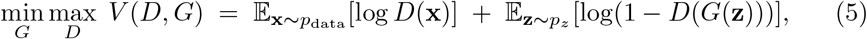

where *p*_data_ is the real data distribution and *p*_*z*_ is a simple prior (e.g., 𝒩 (0, *I*)).

- *Attention-weighted seed pooling:* Let *π*_*i*_ be the priority score for node *i*; let 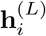 be the final GAT embedding of node *i*. We form an attention-pooled seed from the top-*q*% set *S*_*q*_:

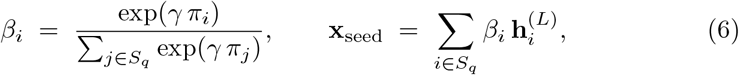

where *γ* ≥ 0 controls sharpness. Optionally, **x**_seed_ is mapped to a generator “conditional” channel by 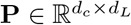.
- *Hybrid generator input (mixed input):* For a fresh noise draw **z** ∼ *p*_*z*_, we build the hybrid input by concatenation:

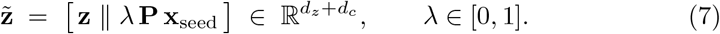

The scalar *λ* is the leakage rate that scales how strongly the seed steers the generator. (Equivalently, a Bernoulli gate *b* ∼ Bernoulli(*λ*) can be used to stochastically toggle the seed channel.)

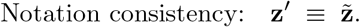
- *GARAGE objective:* Substituting the hybrid input for the generator yields

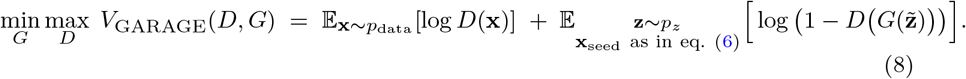

This is equivalent to writing the expectation over the induced hybrid input law 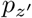 with 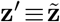.

i. The construction in equations (6)–(7) makes the role of leakage explicit and avoids ambiguity about where to “insert” seeds.
ii. When class labels are available, one can add a weak ratio-matching regularizer 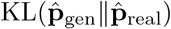 to preserve cell-type proportions, without changing the placement of equation (7).
iii. In our convergence analysis (section 3.6), equation (7) is the reference equation used to derive variance reduction and divergence contraction.

The detailed algorithm for the entire *GARAGE* framework can be found in the algorithm 1.

### 3.6 Convergence properties of *GARAGE*

We provide a theoretical account for why the *GARAGE* objective converges faster than a standard GAN objective under mild regularity assumptions. The core mechanism is that attention-guided leakage induces (i) a variance–reduced stochastic gradient for the generator, and (ii) a contraction of divergence to the data distribution through convex mixing with a seed–anchored component that lies near the real manifold. Together, these effects accelerate stochastic min–max training in expectation.

***Setup:*** Let *p*_data_ denote the real data distribution on and let 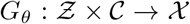 be the generator with parameters *θ*. A standard GAN uses inputs (*z, c*) = (*z*, **0**) with *z ∼ p*_*z*_. In *GARAGE*, the generator sees hybrid inputs

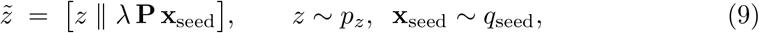

as in equation (7), where *q*_seed_ is the attention–pooled seed distribution in equation (6) and *λ ∈* [0, 1] is the leakage rate. Write 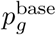 for the pushforward of *p*_*z*_ by *G*_*θ*_(·, **0**), and 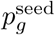 for the pushforward of (*z*, **Px**_seed_) by *G*_*θ*_. Then the *instantaneous* model distribution under leakage is the convex mixture

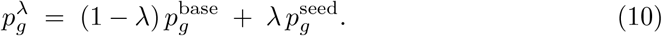

We analyzed the generator update under the non-saturating GAN loss with an optimal–or–near–optimal discriminator *D*_*ϕ*_ per iteration (standard two-time scale assumption).

**Assumption 1**

i. For every (*z, c*), the per–sample generator loss *ℓ*(*θ*; *z, c*) ≡ − log *D*_*ϕ*_ *G*_*θ*_ ([*z*∥*c*]) is L–smooth in θ and has bounded gradient ∥∇_*θ*_*ℓ*(*θ*; *z, c*)∥ ≤ *G*_max_.
ii. *G*_*θ*_ is *L*_in_–Lipschitz in its input and *L*_*θ*_ –Lipschitz in *θ*.
iii. Seeds lie *ε*–close to the real manifold in an integral probability metric (IPM), e.g. *W*_1 (_*q*_seed_, *p*_data)_≤ *ε*.
iv. The discriminator class contains 1–Lipschitz critics and is trained to *δ*–optimality per iteration (small but nonzero).
v. The generator objective satisfies a Polyak–L ojasiewicz (PL) inequality in a neighborhood of a stationary point: 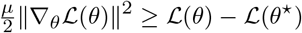 with *µ >* 0.

The assumption 1 follows standard GAN convergence analyses and is reasonable in practice after spectral normalization, gradient penalties, and small steps.

#### Algorithm 1

The GARAGE framework algorithm

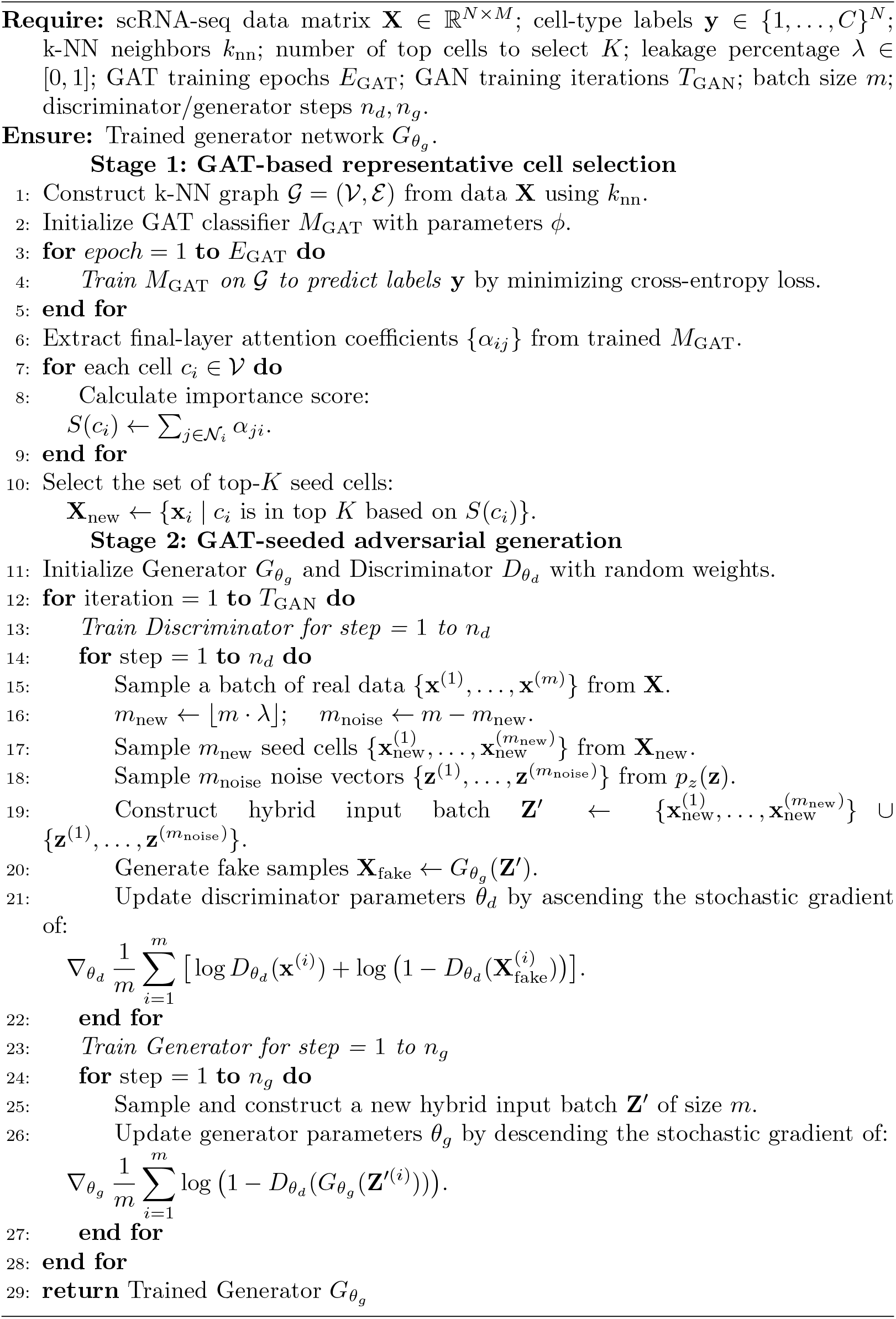

#### 3.6.1 Variance reduction of the generator gradient

Let **g**_base_(*θ*; *z*) = ∇_*θ*_*ℓ*(*θ*; *z*, **0**) and **g**_seed_(*θ*; *z*, **x**_seed_) = ∇_*θ*_*ℓ*(*θ*; *z*, **Px**_seed_). The *GARAGE* stochastic gradient draws

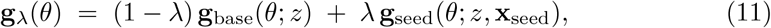

with (*z*, **x**_seed_) independent. Denote the means and variances by ***µ***_b_ = 𝔼[**g**_base_], ***µ***_s_ = 𝔼[**g**_seed_], *Σ*_b_ = Var(**g**_base_), *Σ*_s_ = Var(**g**_seed_).

##### Lemma 1

(Law of total variance for the hybrid gradient) For any *θ*,

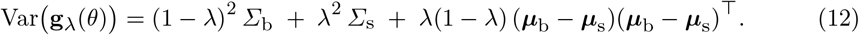

In particular, if *Σ*_s_ ⪯ *Σ*_b_ and ∥***µ***_b_ − ***µ***_s_∥ is bounded, then for all *λ* ∈ (0, 1],

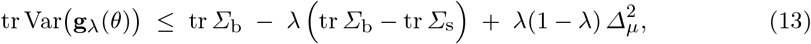

where *Δ*_*µ*_ = ∥***µ***_b_ − ***µ***_s_∥.

Attention–guided seeds concentrate *G*_*θ*_’s input on high–signal regions, which empirically yields *Σ*_s_ ≪ *Σ*_b_. The equation (13) shows that the stochastic gradient variance is reduced linearly in *λ*, up to a small mean–mismatch penalty 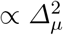. Variance reduction tightens the noise floor of SGD, thereby decreasing the iteration complexity.

##### Proposition 1

(Iteration complexity under PL with variance reduction) *Under the assumption 1 and with a constant step size η* ≤ 1*/L, the generator iterates of GARAGE using mini–batches of size B satisfy*

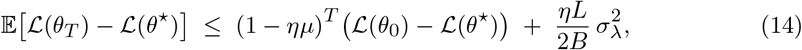

*where* 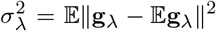 *and, by lemma 1*, 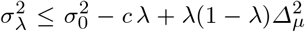 *for some c > 0. Thus for any target accuracy* 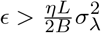, the number of iterations *T*_*λ*_*needed by GARAGE to reach ϵ satisfies* 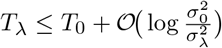 *and in particular T*_λ_ < *T*_0_ whenever 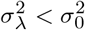.

#### 3.6.2 Divergence contraction by convex mixing with seed–anchored generator

Let D_*f*_ (*·*∥*·*) be any *f* –divergence. It is convex in its second argument, hence for any distributions *q, r* and *λ ∈* [0, 1],

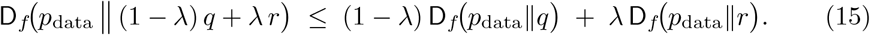

Applying the equation (15) to 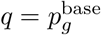 and 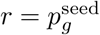 we obtain:

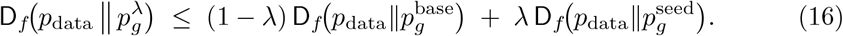

Under the assumption 1(iii)–(iv), the seed–anchored generator is close to the data in an IPM (and hence in common *f* –divergences up to constants), i.e. 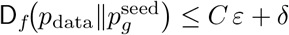. Therefore,

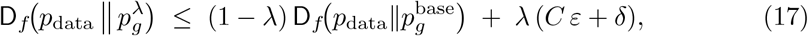

which contracts the divergence to the data by a factor (1 *− λ*) up to a small bias term.

##### Lemma 2

(Avoiding discriminator saturation:) In the non–saturating GAN, the generator gradient magnitude is proportional to 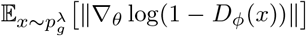. If *D* is a 1–Lipschitz critic close to the IPM–optimal discriminator, then

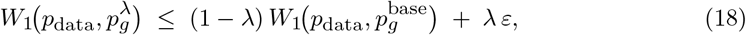

and discriminator logits remain in a non–saturated band that is wider than in the base case, yielding a uniform lower bound on generator gradient norms and improved conditioning.

#### 3.6.3 GARAGE exhibits faster convergence

##### Theorem 1

(Convergence acceleration with attention–guided leakage) *Under the assumption 1, for any λ* ∈ [0, 1] *such that* 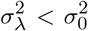 *and with discriminator trained to δ–optimality per iteration, the generator iterates of GARAGE with step size η* ≤ 1*/L satisfy*

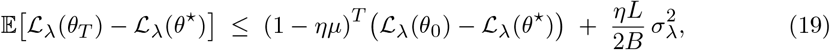

*with* ℒ_*λ*_ *the GARAGE generator objective induced by* 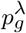. *Moreover, by equation* (17) *and lemma 2, the bias term in equation* (19) *corresponds to a discriminator that is less saturated than in the base GAN, so that for any target accuracy ϵ above the noise floor, the iteration complexity T*_*λ*_ *satisfies T*_*λ*_ *< T*_0_ *whenever λ >* 0 *is chosen so that* 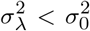 *and C ε* + *δ is small*.

***Proof:*** The equation (12) follows by direct expansion and independence of (*z*, **x**_seed_). The PL–based bound equation (14) is standard for smooth nonconvex objectives under SGD with bounded variance; replacing 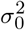 by 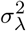 yields equation (14). The convexity of *f* –divergences in the second argument gives equation (16), and the proximity of 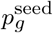 to *p*_data_ (assump-tion 1iii–iv) yields equation (17). For lemma 2, the IPM bound implies discriminator outputs stay away from the saturated regimes, preserving generator gradients. Combining the variance reduction (lower noise floor) with improved conditioning provides the equation (19) and strictly fewer iterations for any fixed *ϵ* above the floor. ? □

#### 3.6.4 Choice of leakage rate

From equation (12), the scalar variance proxy satisfies

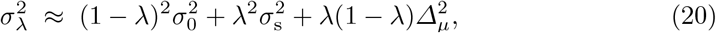

which is minimized near

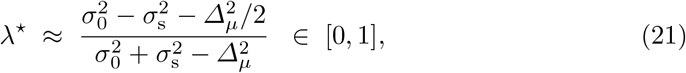

projected to [0, 1]. In practice, a moderate *λ* balances variance reduction against mean–mismatch and avoids over–constraining *G*_*θ*_, consistent with our sensitivity studies.

## 4 Conclusion

In this work, we introduced *GARAGE*, a novel two-stage generative framework designed to address the significant challenge of generating high-fidelity synthetic single-cell RNA-seq data. We identified a key limitation in traditional generative adversarial networks (GANs) - their potential for training instability and the generation of biologically implausible cell states when learning from a random noise distribution alone. To overcome this, GARAGE uniquely integrates a graph attention network (GAT) as an intelligent selection module to identify biologically representative “seed” cells, which are then used to guide and anchor the generative process of a subsequent GAN. Our comprehensive experimental evaluation demonstrates the clear superiority of the GARAGE framework across multiple datasets and performance metrics. Quantitative assessment using the Wasserstein distance consistently revealed that all variants of GARAGE (with 10-30% GAT-informed data leakage) generated distributions significantly closer to the real data than a traditional GAN (0% leakage) across all four benchmark datasets. Furthermore, in a rigorous benchmarking study against leading generative models, including VAE, FGAN, and WGAN, GARAGE consistently achieved state-of-the-art results in downstream clustering fidelity, as measured by the Adjusted Rand Index (ARI), Normalized Mutual Information (NMI), and macro-F1 score. These results affirm that the synthetic data produced by GARAGE not only mirrors the statistical properties of the real data but also crucially preserves the intricate population structures and cell-type identities necessary for meaningful biological analysis.

The success of the GARAGE framework carries significant implications for the field of bioinformatics. It provides a robust method for data augmentation, particularly for studies involving rare cell types where data scarcity is a major bottleneck. Moreover, it enables the creation of high-quality, privacy-preserving synthetic datasets that can be widely shared for algorithm benchmarking and method development without compromising patient confidentiality. The ability to generate synthetic data that accurately preserves complex, multi-modal population structures is a critical step towards more robust and reproducible single-cell analysis.

While GARAGE has demonstrated significant advantages, we acknowledge several avenues for future research. The initial k-NN graph construction and GAT training can be computationally intensive for extremely large datasets, and future work could explore more scalable graph construction or representation learning techniques. Additionally, our results indicate that The optimal leakage percentage *λ* is dataset-dependent; developing an adaptive or learned approach to determine this hyperparameter could further enhance the model’s performance and usability. Future research could also extend the GARAGE framework to other single-cell modalities, such as scATAC-seq or spatial transcriptomics, where modeling intercellular relationships is equally crucial.

In conclusion, the GARAGE model offers a novel and effective approach to generating synthetic single-cell data. By uniquely fusing the discriminative power of graph attention with the generative capabilities of adversarial networks, GARAGE provides a principled approach to overcoming the limitations of traditional methods, paving the way for more reliable and advanced applications of synthetic data in computational biology.

## 5 Software version

All the analyses are performed using R (v4.5.1), Python (v3.12.5) for the GARAGE implementation, Python (v3.7.12) for the benchmarking of SOTA models, and Python (v3.9.21) for the scRNA-seq preprocessing and *GARAGE* model validation. For the preprocessing of the scRNA-seq data, Scanpy (v1.9.6) and scikit-learn (v1.5.2) were utilised. For *GARAGE* architecture building and implementation, we used Torch (v2.5.1 + cu118) and Torch-Geometric (v2.7.0). For Wasserstein distance calculations, we used SciPy (v1.13.1).

## 6 Acknowledgments

The authors would like to sincerely thank Ms Sruti Dey for her initial support with coding and early drafting of the manuscript. Although these contributions were limited to the preliminary stages of the project, they provided a helpful starting point for the subsequent development of the work. The authors are grateful to Ms Nilanjana Bhattacharya for her kind encouragement and for the limited writing assistance she provided during the early stages of this work.

## 7 Author contributions

S.R. and M.H. took the lead in formulating the study concept, designing the methodology, and guiding the overall research framework. Together, R.G. and S.A. developed the programming codes, carried out the data analysis, and produced the first draft of the manuscript. S.R. and M.H. contributed substantial intellectual feedback and played a crucial role in revising the manuscript. All authors carefully reviewed the complete work and approved the final version for submission.

## Notes

### Competing Interest Statement

The authors have declared no competing interest.

